# TX2P: A Proteogenomic Tool for Comprehensive Transcript Analysis

**DOI:** 10.1101/2025.06.24.661260

**Authors:** David Murphy, Mina Ryten, Nicholas W. Wood, Emil K. Gustavsson

## Abstract

Long-read RNA sequencing has expanded our understanding of the transcriptome, revealing unannotated transcripts. However, interpreting their functional relevance remains challenging. To address this, we developed TX2P, a user-friendly tool that integrates transcriptomic and proteomic data to link RNA discoveries with protein function. Applied to epilepsy-associated genes, TX2P identified novel protein-coding transcripts, with peptide evidence supporting 17.2% transcripts and 18.0% of unique open reading frames. By seamlessly integrating transcriptomic and proteomic insights, TX2P facilitates transcriptome interpretation.

## MAIN TEXT

The vast majority of mammalian genes undergo alternative splicing, leading to the production of multiple distinct transcript structures^1^. However, detecting and quantifying these transcripts across different cell types, tissues, and species has been a major challenge. This is primarily due to the significant length disparity between the transcripts and the short reads commonly used in traditional RNA-seq approaches^2^. The emergence of long-read sequencing technologies enables capture of entire RNA molecules within a single read, offering more precise annotation and quantification of transcripts, including the identification of *de novo* structures. Recent studies have demonstrated that long-read sequencing can discover a large fraction of novel transcripts, including those predicted to encode novel protein isoforms, both transcriptome-wide^3–6^ and surprisingly, also for extensively studied disease genes^7,8^.

The increasing availability of long-read RNA-seq datasets highlights the importance of accurate transcript annotation, particularly in determining whether transcripts are translated and can produce alternative protein isoforms. Combining sample-matched long-read RNA-seq with mass spectrometry (MS)-based proteomics has proven effective for isoform characterization^9^. However, this approach poses significant technical and methodological challenges. Besides the requirement for advanced sequencing technologies and high-resolution MS platforms, there is an urgent need for computational tools enabling accurate integration and analysis of transcriptomic and proteomic data. To address this challenge, we developed Transcript-To-Protein (TX2P), a versatile tool for assessing the protein-coding potential of novel transcripts. TX2P predicts open reading frames (ORFs) from transcript sequences and integrates these predictions with mass spectrometry (MS) data, supporting both raw and publicly available MS datasets. By enabling seamless transcriptomic and –proteomic data integration without reliance on proprietary data or software, TX2P streamlines the interpretation of novel transcript discovery and facilitates the identification of protein-coding isoforms.

The TX2P workflow consists of two modules. The first module requires the input of transcript structures formatted in standard GTF/GFF file formats. This is a standard output of a long-read RNA-seq analysis, generated through tools such as Bambu^10^, FLAIR^11^, IsoQuant^12^, StringTie2^13^, and TALON^14^. This module then predicts ORFs and generates the predicted peptide sequences for each transcript. The second module requires raw proteomic datasets, typically from The ProteomeXchange Consortium^15^ and uses MetaMorpheus^16^ for peptide identification. This module also allows for removal of peptide sequences in a reference dataset such as UniProt^17^ to enable the user to focus on novel isoforms alone. A schematic of the workflow is described in **Fig. 1a**.

**Figure 1.**
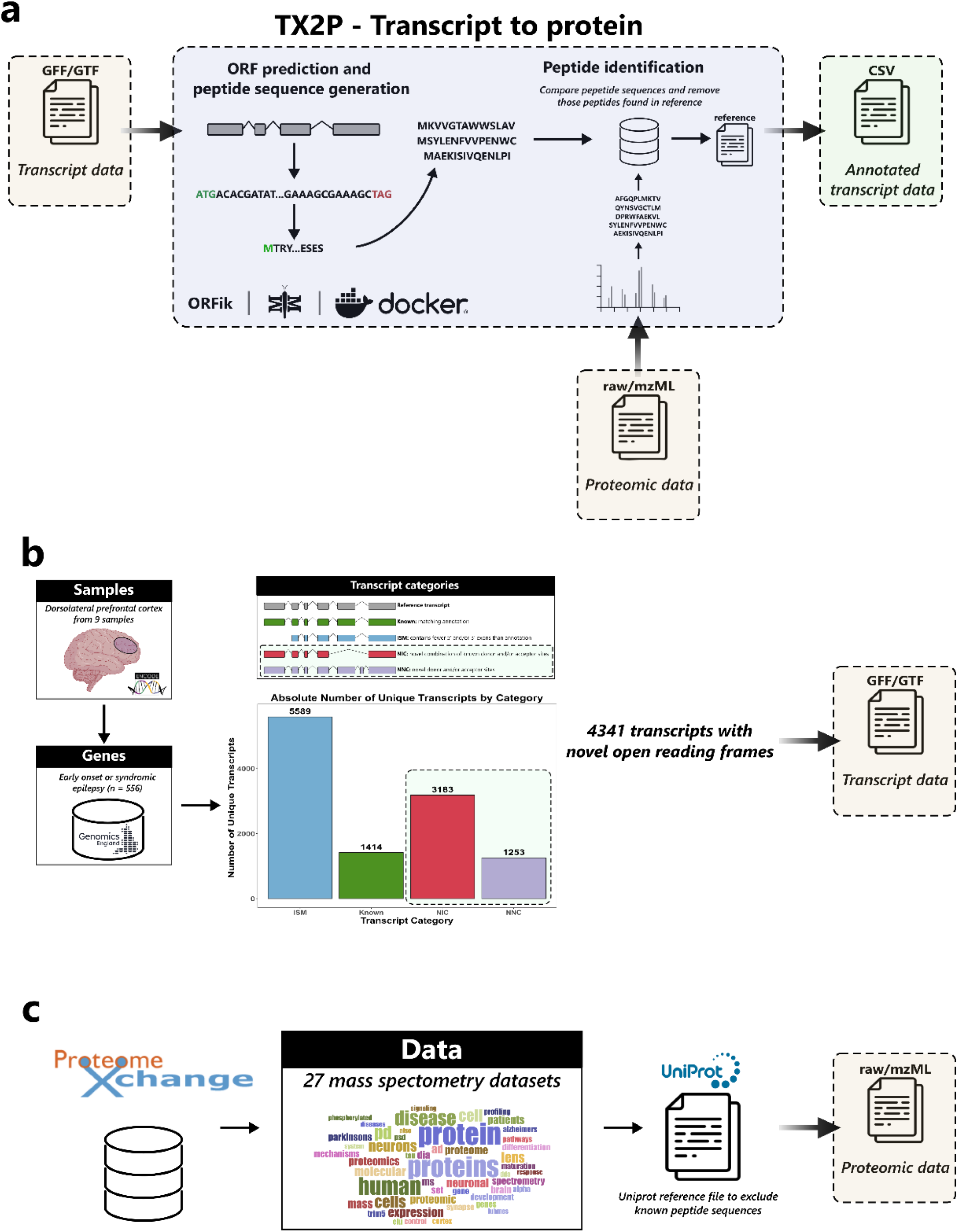
Overview of the TX2P workflow. **(a)** Schematic representation of the TX2P tool, illustrating the key steps in the transcript-to-protein mapping process. **(b)** Methodological framework for generating the input GTF file, detailing how transcript structures were obtained from long-read RNA sequencing data available through the ENCODE project. The analysis focused on novel non-coding (NNC) and novel in catalog (NIC) transcripts. Genes implicated in epilepsy were identified using the Genomics England PanelApp. **(c)** Integration of 27 mass spectrometry datasets from the ProteomeXchange consortium to validate protein-level evidence for transcript-derived open reading frames.

To demonstrate the utility of TX2P, we analysed transcript structures derived from long-read sequencing of human brain samples and leveraged brain-relevant mass spectrometry datasets to find evidence for the translation of novel open reading frames (ORFs) in disease-relevant genes. We focused on 556 unique genes causally implicated in early onset and syndromic epilepsy, sourced from the Genomics England PanelApp^18^ (**Supplementary table 1**). Then, using public long-read RNA-seq data from 9 frontal cortex samples (**Supplementary table 2**), we assessed gene and transcript expression. This data, generated using PacBio Iso-Seq with >99% base pair accuracy enabled by circular consensus sequencing (CCS), had a mean sequencing depth of 2.2 ± 0.9 million full-length reads per sample.

We found that 481 of the 556 genes disease-relevant genes (86.5%) were expressed across all 9 samples, while 36 genes (6.5%) were not expressed in any sample (**Supplementary Fig. 1**). Focusing on the 481 expressed genes, we aimed to identify transcripts with novel open reading frames (ORFs) that arose either through novel splice junctions (NNC) or novel combinations of known splice junctions (NIC), as defined in the Methods section (**Fig. 1b**). This was achieved by merging all NNC and NIC transcript types across all the nine samples using exact coordinate matching. To reduce artefacts, we only included those transcripts supported by ≥ 2 full-length reads across samples. At this step, no filtering was applied based on the number of samples in which a transcript was expressed. Using this approach, a median of 768 ± 329 transcripts per sample was identified, with a total of 4,341 unique transcripts detected (**Supplementary Fig. 2 and Supplementary Table 3**).

The merged GFF file, containing 4,341 transcripts, was processed using TX2P. The first step of TX2P is to predict ORFs, which resulted in a total of 1,978 unique ORFs across these transcripts. Analysis of 27 mass spectrometry datasets identified 514 peptides unique to the predicted amino acid sequences derived from these ORFs, and which did not match those in UniProt (Reviewed dataset UP000005640, dated 2023/09/12). These peptides had a median length of 19 amino acids, range 7 to 59. Furthermore, these 514 peptides provided supporting evidence for the translation of 745 out of the 4,341 transcripts (17.2%) with 356 unique ORFs (18.0%) in at least one of the included MS datasets (**Fig. 2a**).

**Figure 2.**
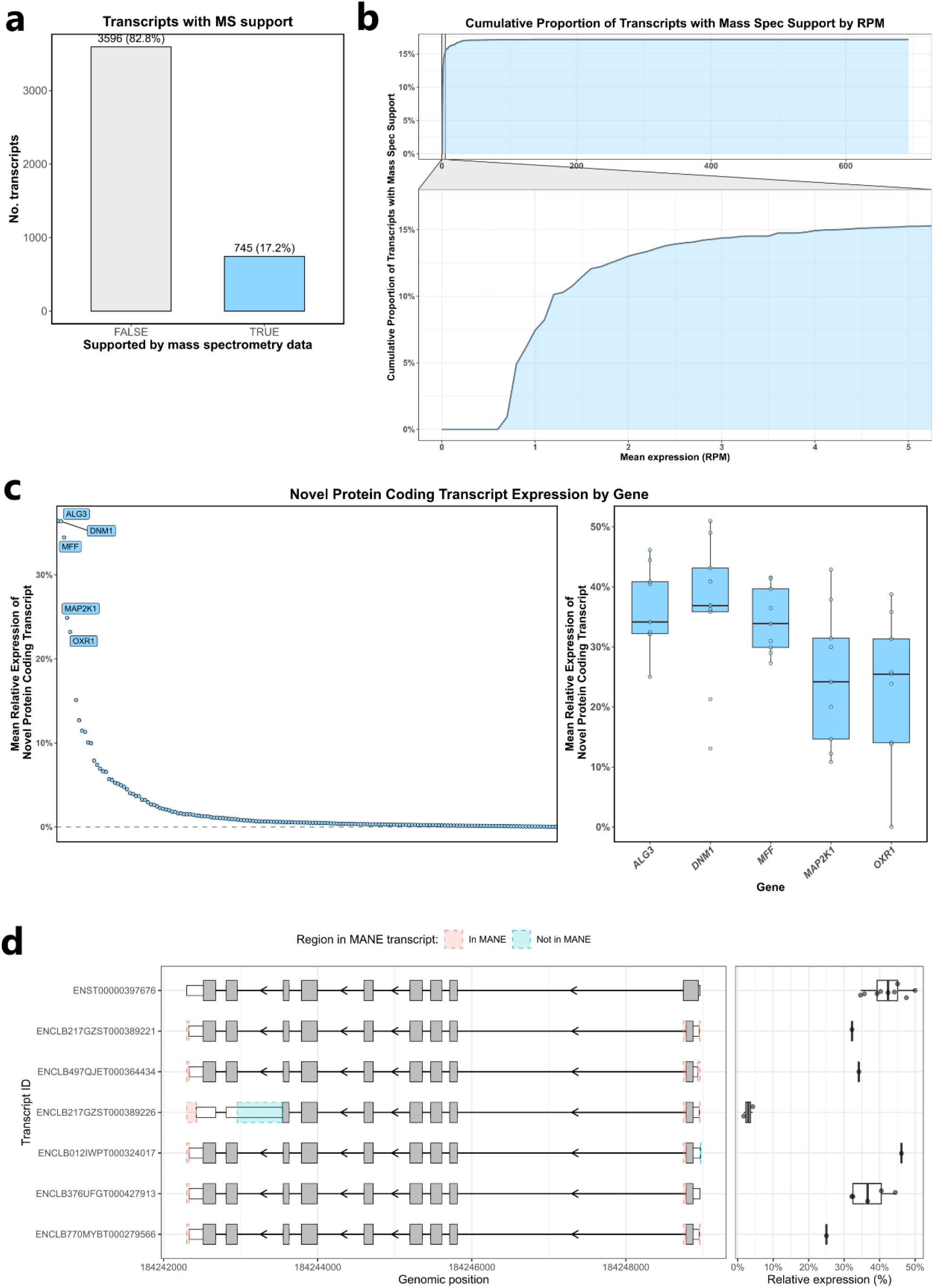
TX2P analysis of expression and proteomic support for novel transcript isoforms in epilepsy-associated genes. **(a)** Total number of transcripts with novel open reading frames (ORF) proteomic support, identified using TX2P. Support was defined by the presence of at least one uniquely mapping peptide in mass spectrometry data, absent from the reviewed UniProt human proteome. **(b)** Cumulative proportion of transcripts with novel ORFs supported by mass spectrometry data, as identified using TX2P. **(c)** Proportion of total gene expression attributable to novel protein-coding transcripts, summarised across all genes with at least one supported novel isoform (left). The five genes with the highest relative expression from supported novel transcripts, *ALG3, DNM1, MFF, MAP2K1* and *OXR1*, are highlighted in the right panel. **(d)** Transcript structures for *ALG3*, showing the use of a novel splice donor site that shortens the 3′ end of exon 1, resulting in a shorter ORF. The novel isoform is shown in comparison to the MANE Select transcript (ENST00000397676).

While this analysis employed a permissive approach with minimal filtering applied prior to transcript inclusion, we also examined the impact of transcript expression levels. As expected, evidence for translation was improved with increasing expression levels (**Fig. 2**). Notably, our analysis relied on bottom-up mass spectrometry datasets, which identify transcripts based on short peptide sequences. While the inclusion of top-down mass spectrometry datasets would have been ideal, their limited availability in public repositories constrained our analysis.

While our proteogenomic analysis revealed that a substantial proportion of novel transcripts were supported by mass spectrometry data, it remained unclear how much these isoforms contributed to overall gene expression. To address this, we next quantified the expression of the novel protein-coding isoforms on a per-gene basis. Although we initially hypothesised that these transcripts would represent only a minor fraction of total gene expression, this was not consistently the case. In several instances, novel protein coding transcripts accounted for a substantial proportion of the total transcript output from a given locus (**Fig. 2C**). For example, in the gene *ALG3*, novel protein-coding transcripts contributed a mean of 36.0 ± 6.9% of total expression from each loci in the human frontal cortex (**Fig. 2C**). This his was driven by transcripts isoform with a shorter ORF, resulting from the use of a novel splice donor site that shortens the 3′ end of exon 1 (**Fig. 2D**).

These findings have important implications. All genes included in this study are causally implicated in human disease, and their current annotations serve as the foundation for diagnostic testing. Misannotation or incomplete annotation could therefore overlook clinically relevant isoforms or misannotate variants that affect splicing. Furthermore, many of these genes belong to the druggable genome; KCNQ2 & STXBP1, are currently included in gene therapy trials listed on ClinicalTrials.gov. The identification of abundant, previously unannotated isoforms in such genes underscores the need to refine transcript annotations to improve both diagnostic and therapeutic strategies.

Thus, the comprehensive characterisation of the transcriptome is crucial for our understanding of biology, the pathogenesis of human disease, and the design of effective therapies. The advent of long-read sequencing, which enables the sequencing of full-length transcripts, has significantly enhanced our comprehension of transcriptome dynamics and expression patterns. Nevertheless, the validation and annotation of frequently occurring novel transcription have emerged as increasingly essential components in the analysis of long-read RNA sequencing pipelines. One important aspect of this is predicting whether novel transcripts are protein-coding. Therefore, TX2P, which seamlessly integrates long-read RNA sequencing and proteomic data, represents an important advance. To our knowledge, it is the first tool that directly annotates proteomic support from any raw mass spectrometry dataset using a GTF/GFF—the standard output of long-read RNA sequencing. By enabling systematic validation of protein-coding potential at scale, TX2P fills a critical gap in the long-read transcriptomics workflow and empowers researchers to move from transcript discovery to functional interpretation.

## ONLINE METHODS

### Usage

Use of TX2P, requires the input of a list of transcript IDs and a GTF/GFF file with transcript structures. The user is then required to: i) set up the Docker environment by installing Docker and pulling the necessary images, ii) run the ORF prediction to identify open reading frames within the transcripts, iii) download the mass spectrometry datasets of interest and process them with MetaMorpheus within the Docker environment. Finally, the user analyses the output to assess the predicted protein-coding transcripts.

### Defining a list of genes associated with epilepsy

To identify genes known to be causally-implicated in early onset or syndromic epilepsy we used Genomics England PanelApp, a publicly available crowdsourced tool to standardise gene panels^18^. We extracted ‘green’, diagnostic-grade genes from PanelApp considering the Early onset or syndromic epilepsy (Version 4.84) gene panel alone.

### Long-read RNA-sequencing data

To identify full-length transcripts with at least one novel splice junction we used publicly available long-read RNA-seq data provided by ENCODE^19^ (https://www.encodeproject.org/rna-seq/long-read-rna-seq/). We specifically used data generated from human frontal cortex samples (*n* = 9). A description of the samples can be found in **Supplementary Table 2**. All samples were sequenced on the PacBio Sequel II platform and processed with the ENCODE DCC deployment of the TALON pipeline (v2.0.0; https://github.com/ENCODE-DCC/long-read-rna-pipeline)^14^. Novel ORFs were defined as transcripts predicted to have ORFs not present in GENCODE v29.

### Mass spectrometry data

We selected 27 bottom-up mass spectrometry datasets from Proteome Exchange, specifically focusing on human brain tissues. Our inclusion criteria were guided by keywords related to brain physiology, including ‘spinal’, ‘pituitary’, ‘dendritic’, ‘cranial’, ‘cortical’, ‘microglia’, ‘basal’, ‘neuron’, ‘neural’, ‘astrocyte’, ‘cortex’, ‘nervous’, and ‘brain’. To ensure a focus on normal cellular functions, we excluded datasets containing keywords such as ‘cancer’, ‘tumor’, and ‘hela’.

To ensure the accuracy we used the Contaminants.fasta file from MaxQuant, which is typically used in filtering out common laboratory contaminants. We also used the full reviewed human proteome dataset from UniProt (UP000005640, dated 2023/09/12).

## CODE AVAILABILITY

TX2P is accessible at https://github.com/MurphyDavid/TX2P (https://doi.org/10.5281/zenodo.14217928). All the code and description of the analysis steps to retrieve the PanelApp gene list, ENCODE long-read RNA-se data and processing of this data is available through: https://github.com/egustavsson/TX2Protein.git (https://doi.org/10.5281/zenodo.15728507).

Docker Images: murphydaviducl/getorf, Murphydaviducl/metamorpheusdocker (https://doi.org/10.5281/zenodo.14218080)

## DATA AVAILABILITY

No new primary data were collected in this study. All data included here is publicly available. Gene panel: https://panelapp.genomicsengland.co.uk/panels/402/, long-read RNA sequencing data: https://www.encodeproject.org/rna-seq/long-read-rna-seq/ (accession ids in **supplementary table 1**), mass spectrometry data: http://www.proteomexchange.org/ with accession ids MSV000079437, MSV000081480, MSV000084521, MSV000085211, MSV000085305, MSV000085434, MSV000085698, MSV000086944, MSV000087506, MSV000088006, MSV000088094, MSV000088256, MSV000088430, MSV000088660, MSV000088744, MSV000089427, MSV000090711, MSV000091027, MSV000091165, MSV000091887, MSV000091888, PXD004010, PXD006122, PXD015239, PXD024748, PXD024998, PXD026370.

## AVAILABILITY OF SOURCE CODE AND REQUIREMENTS

Project name: TX2P: Transcript to Protein

Project home page: https://github.com/MurphyDavid/TX2P

Documentation: https://github.com/MurphyDavid/TX2P/blob/main/README.md

Programming language: R, Python3, Bash

Other requirements: docker

