## supplementary figures for "TX2P: A Proteogenomic Tool for Comprehensive Transcript Analysis"

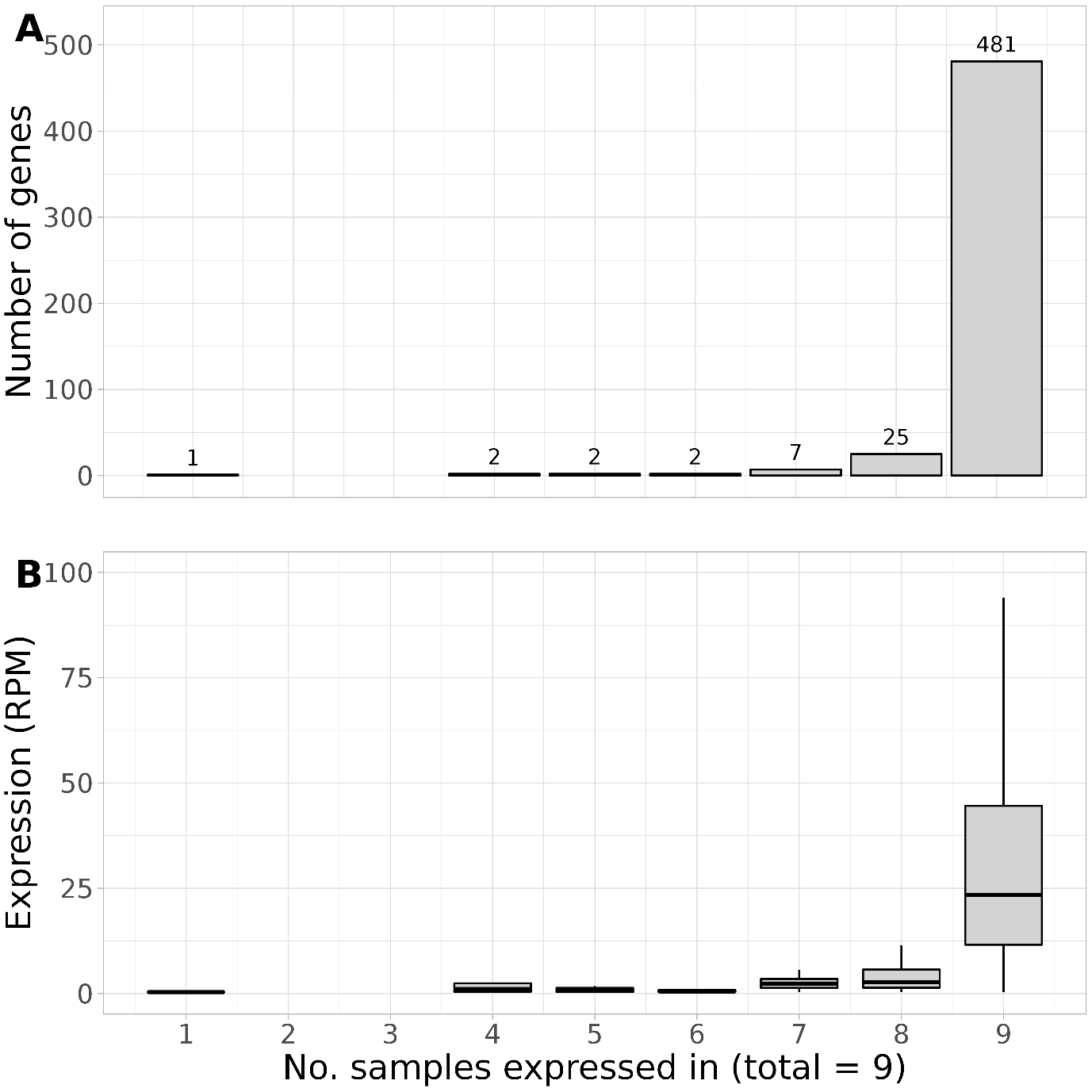

**Supplementary Figure 1. Expression of epilepsy-associated genes in human frontal cortex.** Expression profiles of 556 genes causally implicated in early-onset and syndromic epilepsy, as defined by the Genomics England PanelApp. Long-read RNA-sequencing data from nine human frontal cortex samples (PacBio Iso-Seq) were used to assess gene-level expression. **(a)** Number of samples (out of 9) in which each gene was detected. **(b)** Mean gene expression across samples, measured in reads per million (RPM).

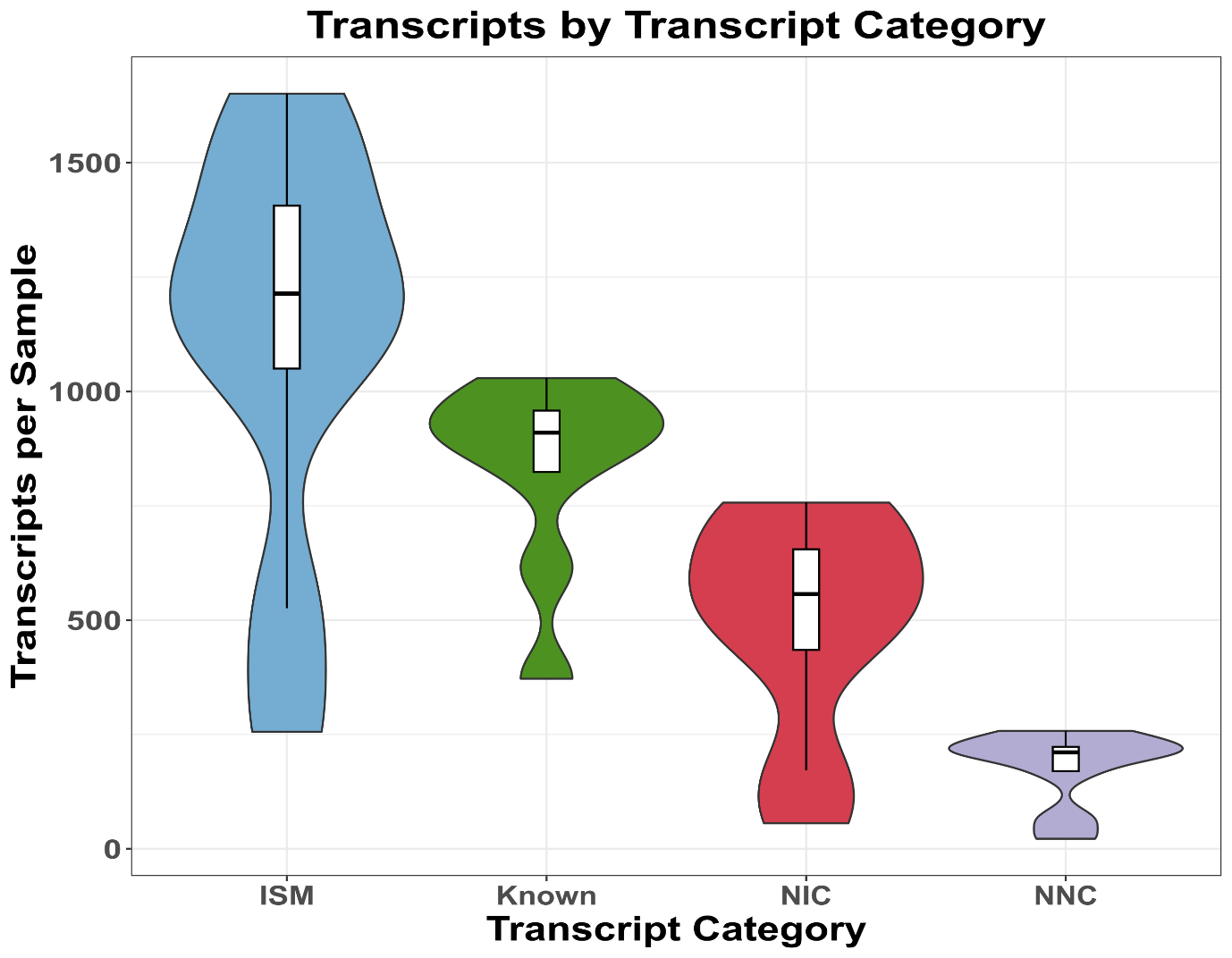

**Supplementary Figure 2. Classification of transcript structures across epilepsy-associated genes.**
Total number of transcripts identified in the long-read RNA-sequencing dataset from nine human frontal cortex samples, grouped by transcript novelty category: Known, Incomplete Splice Match (ISM), Novel In Catalog (NIC), and Novel Not in Catalog (NNC), as defined by the TALON pipeline.

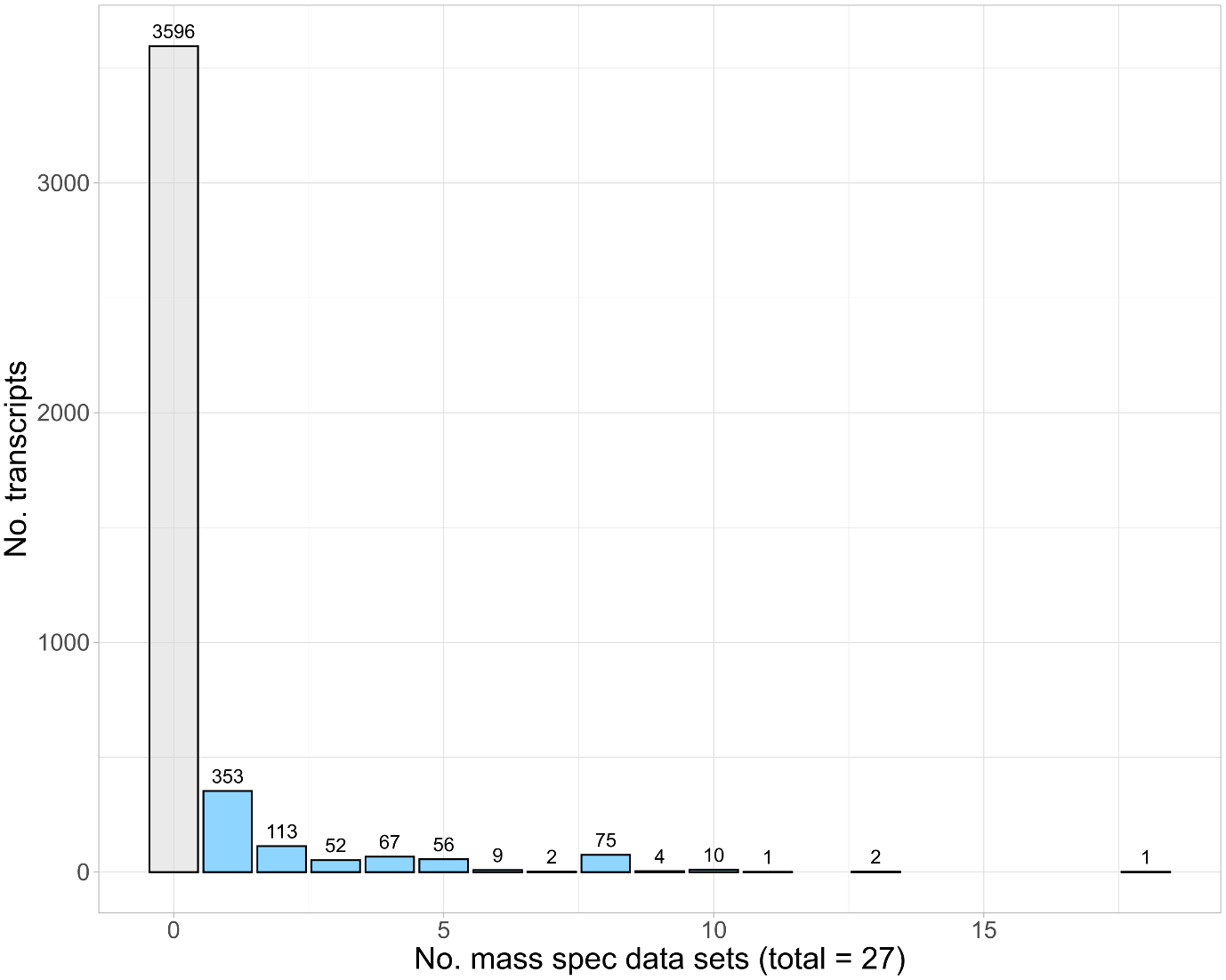

**Supplementary Figure 3. Proteomic support for transcript-derived open reading frames.** Number of transcripts with proteomic support through mass spectrometry, identified using TX2P. Transcripts are grouped by the number of mass spectrometry datasets (out of 27) in which each was supported. Only peptides uniquely mapping to predicted open reading frames and absent from the reviewed UniProt human proteome were considered.
